## Supplementary Figures S1-S11 for "Enterococcal cell wall remodelling underpins pathogenesis via the release of the Enteroccocal Polysaccharide Antigen (EPA)"

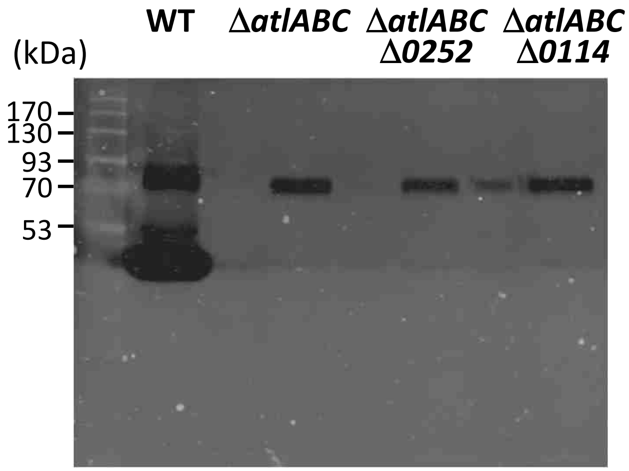


**S1 Fig. Zymogram analysis of *E. faecalis* JH2-2 culture supernatants.** Peptidoglycan hydrolytic activities were detected in 20 µL of culture supernatants of strains JH2-2 (WT), Δ*atlABC,* Δ*atlABC* Δ*0252* and Δ*atlABC* Δ*0114* grown overnight. Cells from the triple class A PBP mutant Δ*ponA* Δ*pbpF* Δ*pbpZ* were used as a substrate and zymograms were incubated for 72h at 37°C.

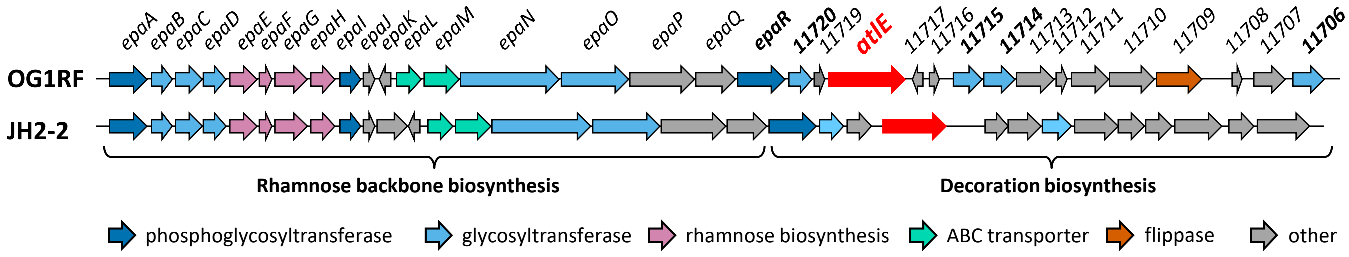


**S2 Fig. Comparison of OG1RF and JH2-2 loci encoding EPA.** Genes *epaA* to *epaR* are conserved across strains *epaR* to 11706 encode EPA decorations which can vary between strains.

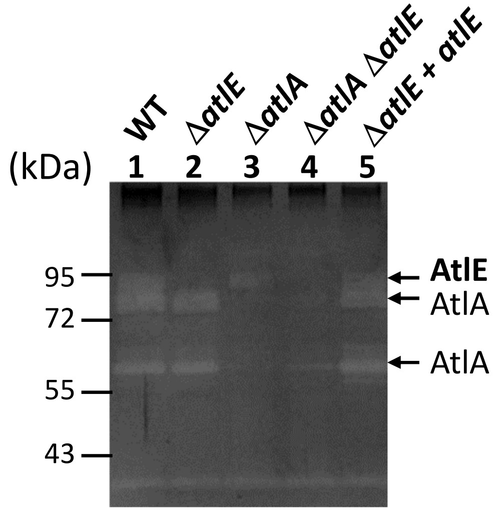


**S3 Fig. Zymogram analysis of *E. faecalis* OG1RF culture supernatants.** Peptidoglycan hydrolytic activities were detected in 25 µL of culture supernatants of strains OG1RF (WT, lane 1), Δ*atlE (lane 2),* Δ*atlA* (lane 3), Δ*atlA* Δ*atlE* (lane 4), and complemented Δ*atlA* Δ*atlE* mutant (Δ*atlE* + *atlE*, lane 5) grown overnight. OG1RF cells were used as a substrate and zymogams were incubated for 72h at 37°C.

A

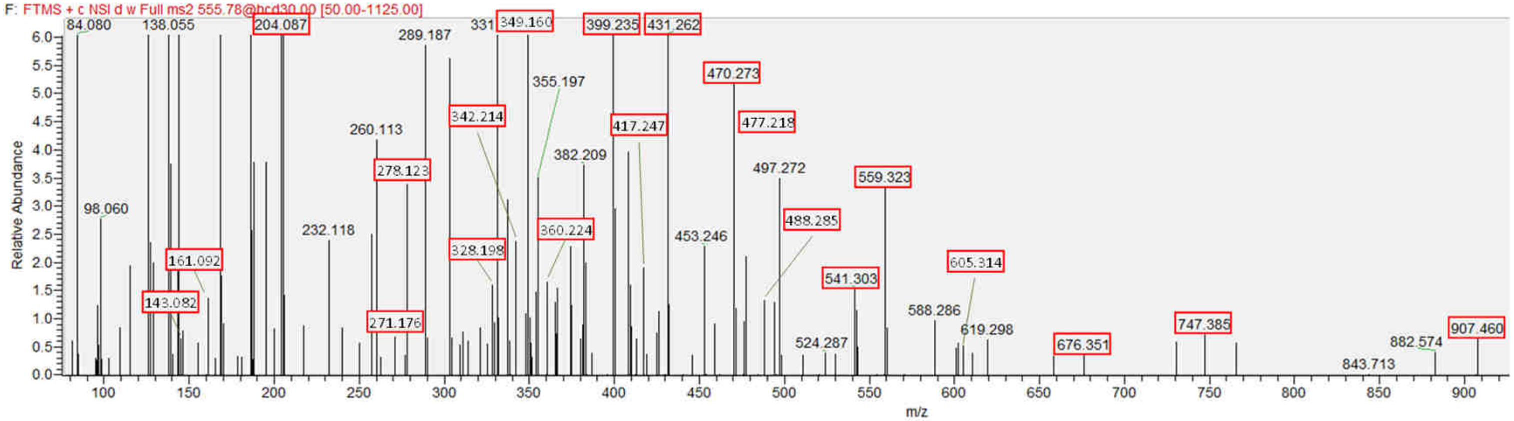


B

| Structure | m/z | | Δppm |
|---|---|---|---|
| | Theo | Obs | |
| gm(r)-AQK[AA]AA | 1110.5525 | ND | |
| gm(r)-AQK[A]AA | 1039.5154 | ND | |
| gm(r)-AQK[AA]A | 1021.5048 | ND | |
| gm(r)-AQKAA | 968.4782 | ND | |
| gm(r)-AQK[A]A, gm(r)-AQK[AA] | 950.4677 | ND | |
| m(r)-AQK[AA]AA | 907.4731 | 907.46 | -14.4 |
| gm(r)-AQK[A], gm(r)-AQKA | 879.4306 | ND | |
| m(r)-AQK[A]AA | 836.4360 | ND | |
| m(r)-AQK[AA]A | 818.4254 | ND | |
| gm(r)-AQK | 808.3935 | ND | |
| m(r)-AQKAA | 765.3989 | ND | |
| m(r)-AQK[A]A, m(r)-AQK[AA] | 747.3883 | 747.385 | -4.4 |
| gm(r)-AQ | 680.2985 | ND | |
| m(r)-AQK[A], m(r)-AQKA | 676.3512 | 676.351 | -0.3 |
| AQK[AA]AA | 630.3570 | ND | |
| m(r)-AQK | 605.3141 | 605.314 | -0.1 |
| QK[AA]AA, AQK[A]AA | 559.3198 | 559.323 | 5.7 |
| gm(r)-A | 552.2399 | ND | |
| AQK[AA]A | 541.3093 | 541.303 | -11.6 |
| QK[A]AA, AQKAA | 488.2827 | 488.285 | 4.7 |
| gm(r) | 481.2028 | ND | |
| m(r)-AQ | 477.2191 | 477.218 | -2.3 |
| QK[AA]A, AQK[A]A, AQK[AA] | 470.2722 | 470.273 | 1.8 |
| K[AA]AA | 431.2613 | 431.262 | 1.7 |
| QKAA | 417.2456 | 417.247 | 3.3 |
| QK[A]A, QK[AA], AQK[A],AQKA | 399.2350 | 399.235 | -0.1 |
| K[A]AA | 360.2241 | 360.224 | -0.4 |
| m(r)-A | 349.1605 | 349.16 | -1.6 |
| K[AA]A | 342.2136 | 342.214 | 1.2 |
| QK[A], AQK, QKA | 328.1979 | 328.198 | 0.2 |
| KAA | 289.1870 | ND | |
| m(r) | 278.1234 | 278.123 | -1.5 |
| K[A]A, K[AA] | 271.1765 | 271.176 | -1.7 |
| QK | 257.1608 | ND | |
| g | 204.0866 | 204.087 | 1.7 |
| K[A], KA | 200.1394 | ND | |
| AQ | 200.1030 | ND | |
| AA | 161.0921 | 161.092 | -0.4 |
| AA | 143.0815 | 143.082 | 3.5 |
| K | 129.1022 | ND | |
| Q | 129.0659 | ND | |
| A (C-ter) | 90.0550 | ND | |
| A (N-ter) | 72.0444 | ND | |

**S4 Fig. MS/MS analysis of peak 5. A**, fragmentation of the doubly charged Ion $(M+2H]^{2+}$; *m/z*=555.78). Ions with *m/z* values matching predicted fragments are boxed in red. **B**, List of predicted fragments, theoretical and observed *m/z*. ND, not detected; g, GlcNAc; m(r), reduced MurNAc; residues in square bracket correspond to the lateral chain.

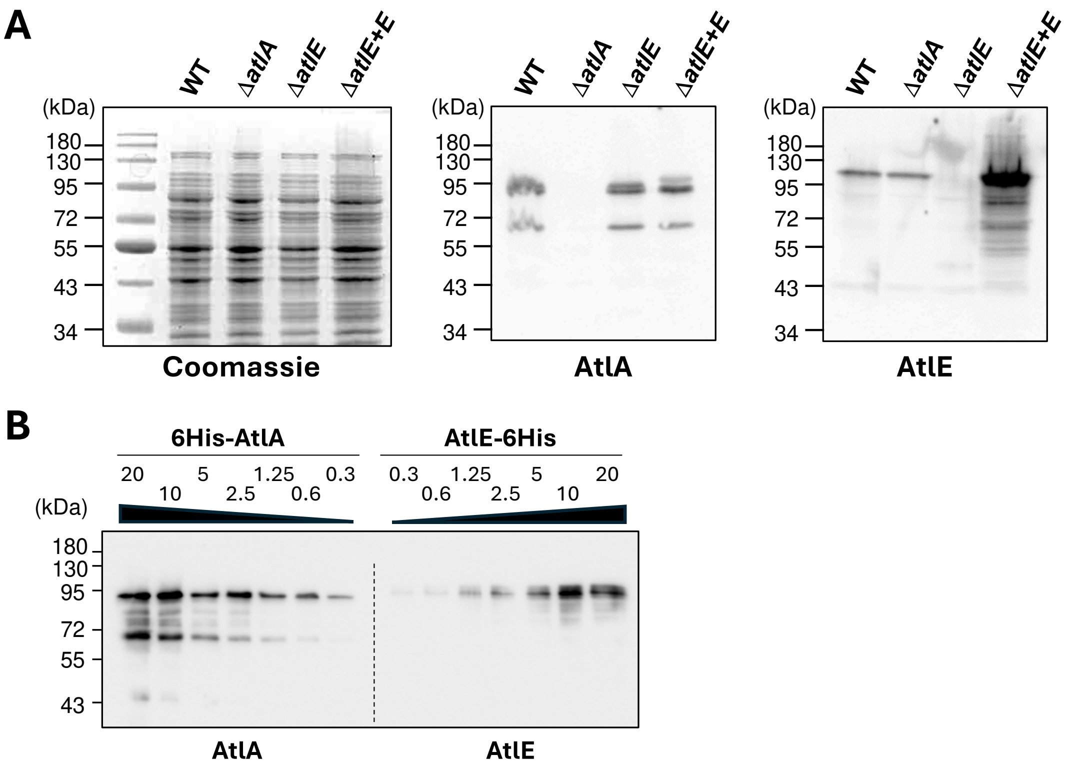


**S5 Fig. Specificity and sensitivity of antibodies recognizing AtlA and AtlE. A,** Specificity of antibodies raised against recombinant AtlA and AtlE proteins was tested against *E. faecalis* crude extracts from cells grown in exponential phase ($OD_{600nm}$≈0.3); WT is OG1RF, Δ*atlE+E* corresponds to the Δ*atlE* mutant complemented. For AtlA detection, 2µg of crude extracts were used; primary serum was used at a dilution of 1/25,000. For AtlE detection, 5µg of crude extracts were used; primary serum was used at a dilution of 1/10,000. In bth cases, secondary antibodies (goat anti-rabbit antibodies coupled to horseradish peroxidase) were used at a 1/20,000 dilution. **B**, sensitivity of anti-AtlA and anti-AtlE antibodies.

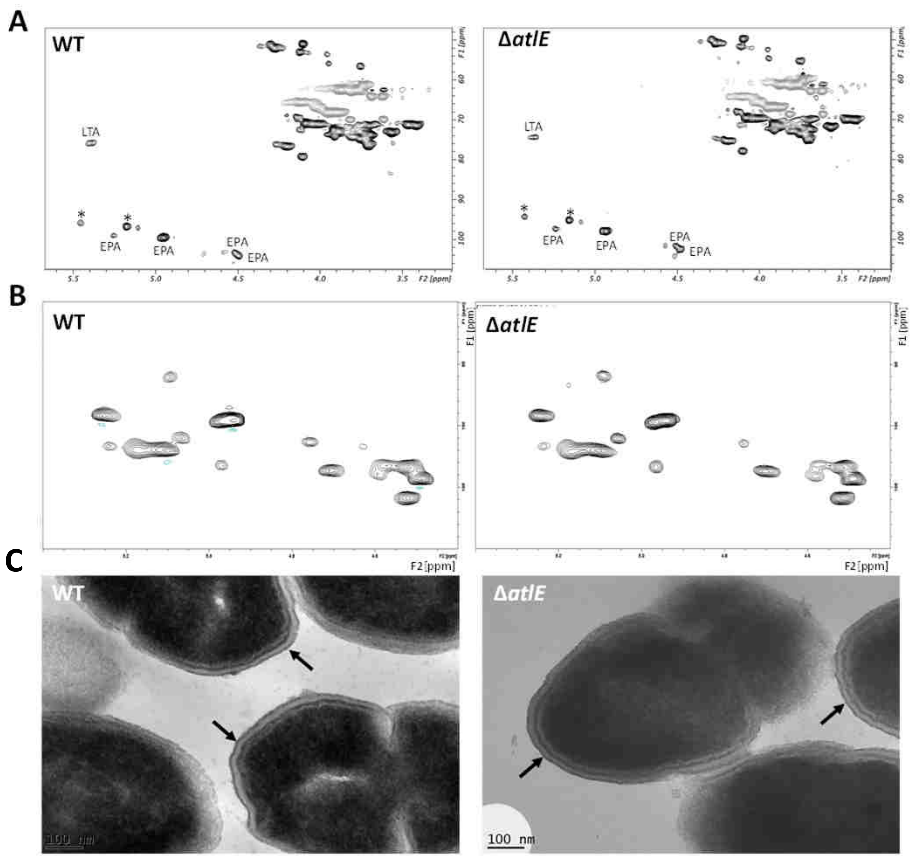


**S6 Fig. AtlE is not involved in EPA biosynthesis or exposure at the cell surface. A,** $^{1}H$-$^{13}C$ HSQC spectra of purified wild type and Δ*atlE* EPA. The region displayed corresponding to the anomeric protons (4.3-5.5 ppm) and anomeric carbons (90-110 ppm) of WT (left) and *ΔatlE* (right) did not reveal any major difference between the 2 EPA polymers. **B**, $^{1}H$-$^{13}C$ HSQC HR-MAS NMR experiments recorded on *E. faecalis* OG1RF (left) and Δ*atlE* (right) cells show that AtlE does not contribute towards the production or display of surface exposed EPA or lipoteichoic acid (LTA) [26]. Two other currently unidentified cell wall polysaccharides denoted with an asterisk are also detected [21]. **C,** Thin section transmission electron microscopy of *E. faecalis* OG1RF (left) and Δ*atlE* (right) cells confirm EPA decorations remain surface exposed in the Δ*atlE* mutant, both forming a pellicle at their cell surface (arrows).

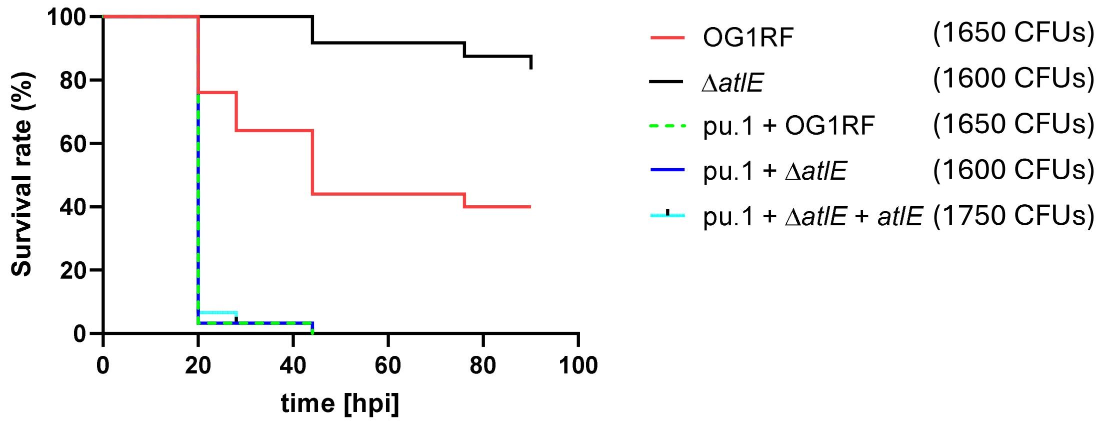


**S7 Fig. Survival rate of zebrafish larvae infected with *E. faecalis* OG1RF, Δ*atlE* and Δ*atlE* complemented strains and role of macrophages in lethality**. Larvae were infected with *ca.* 1600 CFUs of parental (WT) OG1RF strain (red solid line) or Δ*atlE* (black line). Injections were repeated after depletion of macrophages using pu.1 morpholinos; parental (WT) OG1RF strain (green dashed line), Δ*atlE* (blue line), Δ*atlE* + *atlE* (cyan line). Survival was monitored between 20 to 90 hours post infection (hpi) at 28°C using at least 25 larvae per strain per experiment.

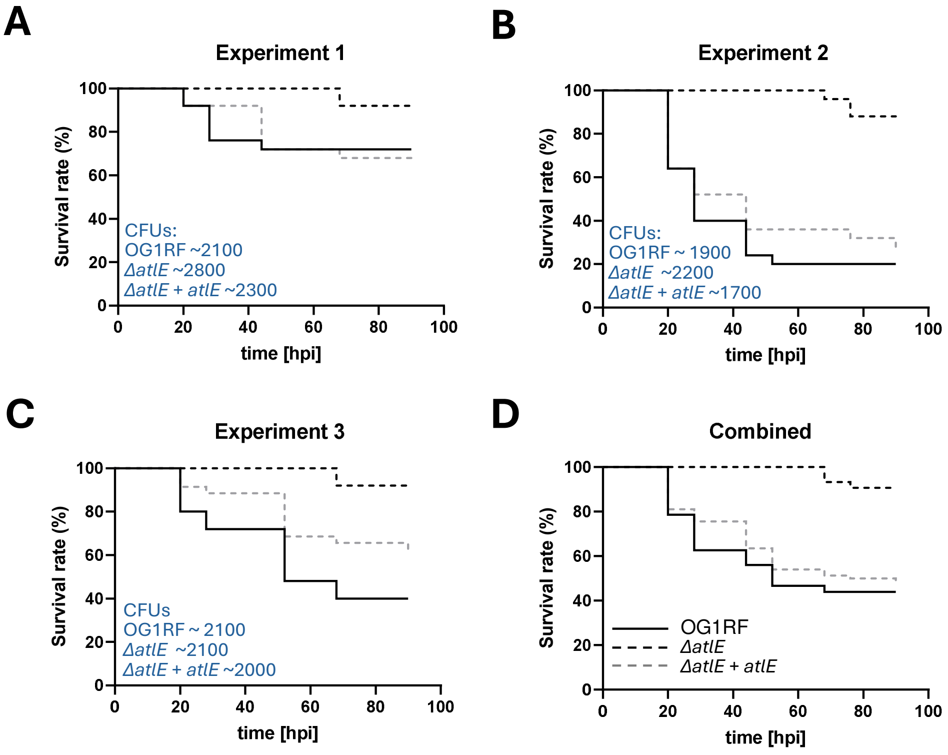


**E**

| | P values | | | |
|---|---|---|---|---|
| | Exp. 1 | Exp. 2 | Exp. 3 | Combined |
| OG1RF vs *Δ11718* | 0.0547 | <0.0001 | <0.0001 | <0.0001 |
| OG1RF vs *Δ11718* pTET2 | 0.9079 | 0.4661 | 0.0587 | 0.4419 |
| *Δ11718* vs *Δ11718* pTET2 | 0.0282 | <0.0001 | 0.0092 | <0.0001 |

**S8 Fig. Survival rate of zebrafish larvae infected with *E. faecalis* OG1RF, Δ*atlE* and Δ*atlE* complemented strains**. Larvae were infected with *ca.* 2000 CFUs of parental (WT) OG1RF strain (solid line), Δ*atlE* (black dashed line) or Δ*atlE* + *atlE* (grey dashed line). Survival was monitored between 20 to 90 hours post infection (hpi) at 28°C using 25 larvae per strain per experiment. Three independent experiments (A, B and C) and combined results (D) are shown. (E) *P* values of pairwise comparison.

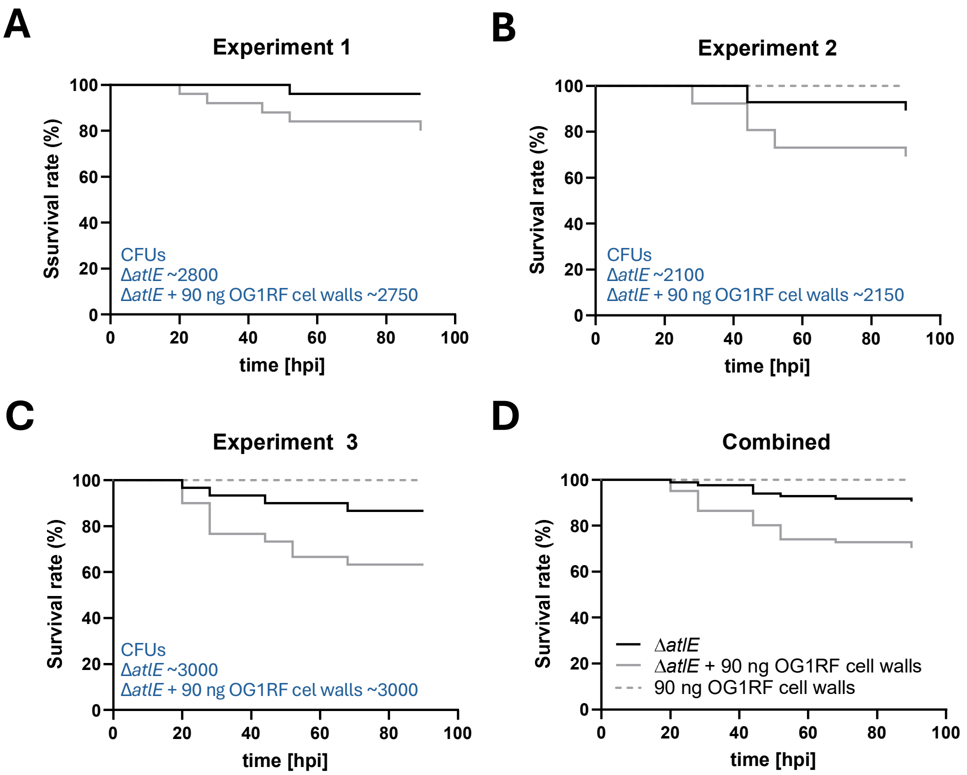


**E**

| | ***P* values** | | | |
|---|---|---|---|---|
| | **Exp. 1** | **Exp. 2** | **Exp. 3** | **Combined** |
| **Δ*atlE* vs Δ*atlE* + OG1RF cell walls** | **0.0724** | **0.0633** | **0.0367** | **0.0010** |

**S9 Fig. Survival rate of zebrafish larvae infected with *E. faecalis* Δ*atlE* in the presence or absence of OG1RF soluble cell wall fragments**. Larvae were infected with *ca.* 2,000 CFUs of the Δ*atlE* strain in the absence (solid line) or presence (grey line) of 90 ng of soluble cell walls. A control injection corresponding to 90 ng of OG1RF cell walls alone is shown (grey dashed line). Survival was monitored between 20 to 90 hours post infection (hpi) at 28°C using at least 25 larvae per strain per experiment. Three independent experiments (A, B and C) and combined results (D) are shown. (E) *P* values of pairwise comparisons.

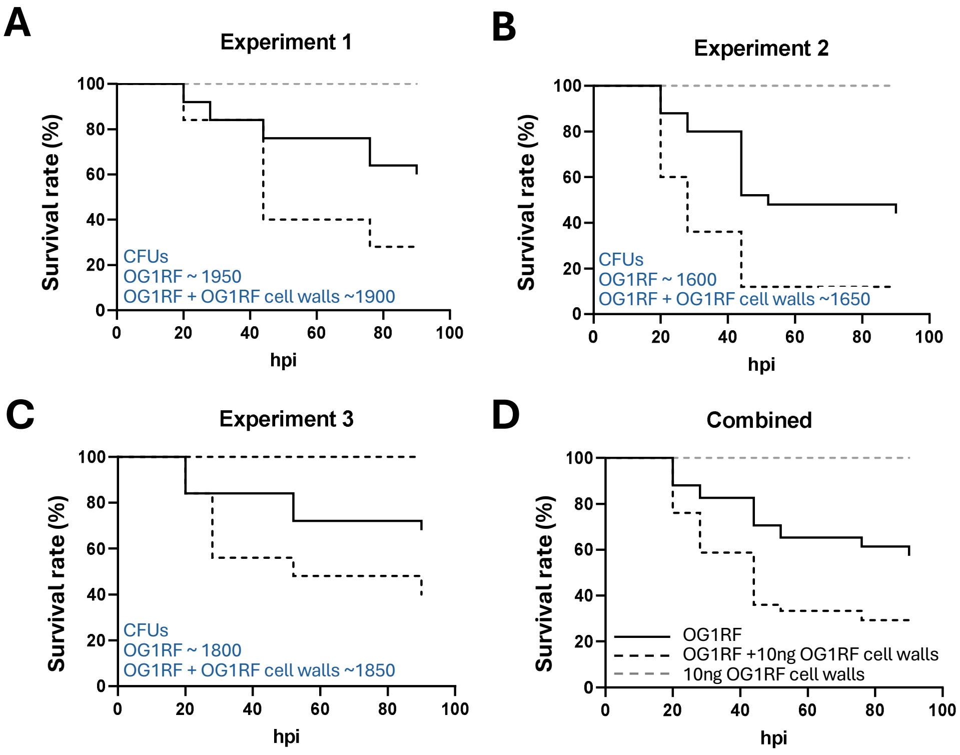


**E**

| | P values | | | |
|---|---|---|---|---|
| | Exp. 1 | Exp. 2 | Exp. 3 | Combined |
| OG1RF vs OG1RF+full EPA | 0.0268 | 0.0013 | 0.053 | <0.0001 |
| OG1RF vs full EPA | 0.0004 | <0.0001 | 0.0022 | <0.0001 |
| OG1RF+full EPA vs full EPA | <0.0001 | <0.0001 | <0.0001 | <0.0001 |

**S10 Fig. Survival rate of zebrafish larvae infected with *E. faecalis* OG1RF in the presence or absence of OG1RF soluble cell wall fragments**. Larvae were infected with *ca.* 2000 CFUs of parental (WT) OG1RF strain in the absence (solid line) or presence (black dashed line) of 10ng of soluble cell walls. A control injection corresponding to 10ng of OG1RF cell walls alone is shown (grey dashed line). Survival was monitored between 20 to 90 hours post infection (hpi) at 28°C using 25 larvae per strain per experiment. Three independent experiments (A, B and C) and combined results (D) are shown. (E) *P* values of pairwise comparison.

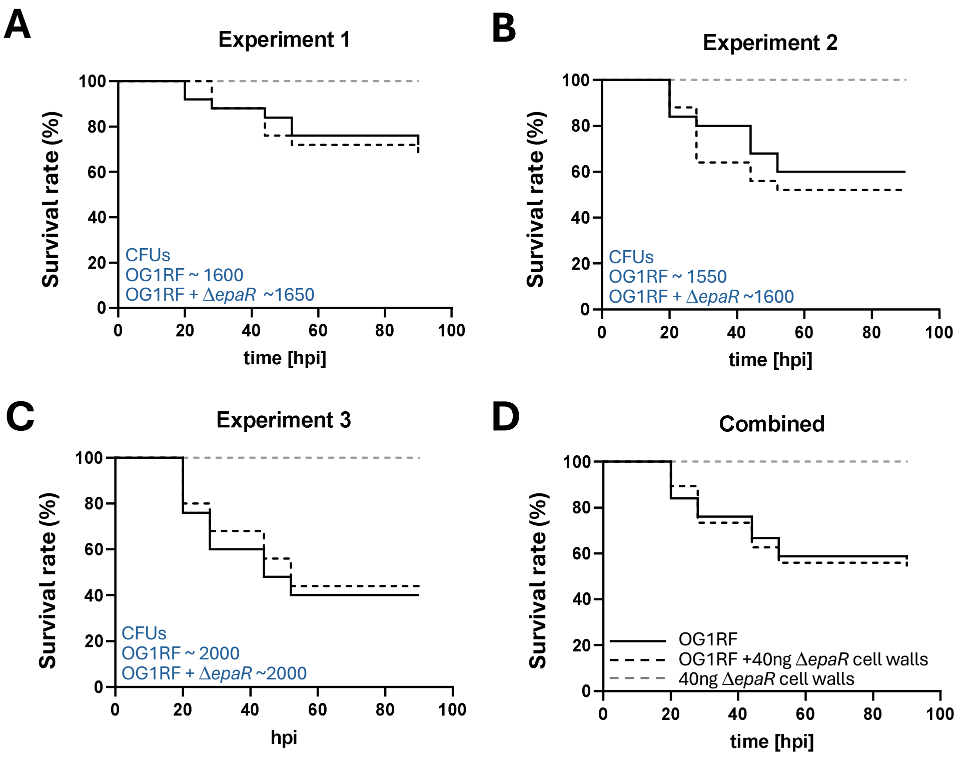


**E**

| | P values | | | |
|---|---|---|---|---|
| | Exp. 1 | Exp. 2 | Exp. 3 | Combined |
| OG1RF vs OG1RF+EPA mutant | 0.7831 | 0.3732 | 0.6912 | 0.7826 |
| OG1RF vs EPA mutant | 0.0047 | 0.0004 | <0.0001 | <0.0001 |
| OG1RF+EPA mutant vs EPA mutant | 0.0022 | <0.0001 | <0.0001 | <0.0001 |

**S11 Fig. Survival rate of zebrafish larvae infected with *E. faecalis* OG1RF in the presence or absence of *epaR* soluble cell wall fragments lacking EPA decorations**. Larvae were infected with *ca.* 2000 CFUs of parental (WT) OG1RF strain in the absence (solid line) or presence (black dashed line) of 40ng of soluble cell walls. A control injection corresponding to 40ng of OG1RF cell walls alone is shown (grey dashed line). Survival was monitored between 20 to 90 hours post infection (hpi) at 28°C using 25 larvae per strain per experiment. Three independent experiments (A, B and C) and combined results (D) are shown. (E) P values of pairwise comparison.
