## Supplementary Table 2 for "Enterococcal cell wall remodelling underpins pathogenesis via the release of the Enteroccocal Polysaccharide Antigen (EPA)"

**Table S1.** Bacterial strains, plasmids, and oligonucleotides.

| Strains/plasmids/ oligonucleotides | Relevant properties/sequence | Source |
|---|---|---|
| **Strains** | | |
| *Enterococcus faecalis* | | |
| JH2-2 | Laboratory strain | (1) |
| JH2-2 Δ*atlABC* | JH2-2 derivative with an in-frame deletions in *atlA*, *atlB* and *atlC* | (2) |
| JH2-2 Δ*atlABC* Δ*0114* | JH2-2 Δ*atlABC* derivative with an in-frame deletions in *TX4000_00147* | This work |
| JH2-2 Δ*atlABC* Δ*0252* | JH2-2 Δ*atlABC* derivative with an in-frame deletions in *TX4000_00259* | This work |
| JH2-2 Δ*atlABC* Δ*atlE* | JH2-2 Δ*atlABC* derivative with an in-frame deletions in *TX4000_01873* | This work |
| JH2-2 Δ*pbp* | JH2-2 Δ*ponA* Δ*pbpF* Δ*pbpZ*; substrate for zymogram experiments | (3) |
| OG1RF | Clinical isolate from human oral cavity | (4) |
| OG1RF Δ*atlA* | OG1RF derivative with an in-frame deletion in *atlA* | (5) |
| OG1RF Δ*atlE* | OG1RF derivative with an in-frame deletion in *OG1RF_11718* | This work |
| OG1RF Δ*atlA* Δ*atlE* | Double *atlA atlE* mutant | This work |
| OG1RF Δ*11720* | OG1RF derivative with an in-frame deletion in *OG1RF_11720* | (6) |
| OG1RF Δ*11720+11720* | OG1RF Δ*11720* complemented strain | (6) |
| *Escherichia coli* | | |
| NEB5alpha | Cloning strain | NEB |
| BL21(DE3) | Protein expression strain | NEB |
| **Plasmids** | | |
| pGhost9 | Plasmid for allelic replacement in *E. faecalis* (Erm$^{R}$) | (7) |
| pTetH | Plasmid for complementation (anhydrotetracyline-induced expression) (Erm$^{R}$) | (5) |
| pET2818 | pET derivative for protein production (C-terminal histidine-tag) (Amp$^{R}$) | (8) |
| pGHH0252 | pGhost9 derivative for *EF0252* deletion | This work |
| pGHH0114 | pGhost9 derivative for *EF0114* deletion | This work |
| pGHH_atlE_J | pGhost9 derivative for *atlE* deletion in JH2-2 | This work |
| pGHH_atlE_O | pGhost9 derivative for *atlE* deletion in OG1RF | This work |
| pET_AtlE_O | pET2818 derivative encoding AtlE allele from OG1RF (residues 25 to 818) (Amp$^{R}$) | This work |
| pTet_AtlE_J | pTetH derivative encoding full length AtlE from JH2-2 for complementation | This work |
| pTet_AtlE_O | pTetH derivative encoding full length AtlE from OG1RF for complementation | This work |
| **Oligonucleotides** | | |
| EF0252_H11 | TATAGGGCGAATTGGGTACCGGGCCCCCCCTCGAGAACCTTTAGAAGGAATTGAACGGAAAATT | |
| EF0252_H12 | GTTATTACCTTCTGCAAATGCACCTACTGGC | |
| EF0252_H21 | GGTGCATTTGCAGAAGGTAATAACAAGGGATTAAACGTTGTTTCGACACGTA | |
| EF0252_H22 | CTCTAGCTAGTGGATCCCCCGGGCTGCAGGAATTCGCGTTTCCCGAAGCGGTTTTCC | |
| EF0252_H110 | TCTATTACGGGCGACAGGGGTCG | |
| EF0252_H220 | GGTCTGCACTGGCAGGGACATCAAT | |
| EF0114_H11 | TATAGGGCGAATTGGGTACCGGGCCCCCCCTCGAGCAGCCAGGCCAGAAAGTCCTGATT | |
| EF0114_H12 | TGCCAAGCCAACCAATAACGAAAGACC | |
| EF0114_H21 | GTCTTTCGTTATTGGTTGGCTTGGCACCAATTATTGGAAGTATTCAATGTGTTTGG | |
| EF0114_H22 | CTCTAGCTAGTGGATCCCCCGGGCTGCAGGAATTCAAGTGTCTCCATTTGAACAGAAGC | |
| EF0114_H110 | GTCGGTTTGACAAGTTATCAAGCGC | |
| EF0114_H220 | CCAGCTTGAGGTGCATTAGGGATAG | |
| atlE_H11_J | AAACTCGAGTTATTAGGGATTTTTTCTTCAAGCAAAATTCGCT | |
| atlE_H12_J | CACTATTGATGAACTAGGAACCAATGCTGT | |
| atlE_H21_J | TTCCTAGTTCATCAATAGTGGCTGGGTAGATAGTCGAGCATTAAAAAAATAAC | |
| atlE_H22_J | CAGGAATTCCCTGTCCATAATACATTCTTTAACAAACCCTACC | |
| atlE_H11_O | AAACTCGAGAACCACCTCTACATCTTCTAACAAATGG | |
| atlE_H12_O | TTTATTTGAAGTGATATACCCTTGTGATGCTTGAGTGACCAAGCCAG | |
| atlE_H21_O | CAACGGGGGAGACTGGCTTGGTCACTCAAGCATCACAAGGGTATATCACTTCA | |
| atlE_H22_O | CAGGAATTCGATATCAAGCTTCATTCACCCCTGCTCTTAC | |
| atlE_H110 | TTTTATGCCACGACCAGATTATG | |
| atlE_H220 | AAATTGTTTAGGAATCTCCTGCC | |
| atlE_J_Fw | CGTGAGCTCAAGGAGGAGACTGACCATGAAGAGAATAAATAAAATATCTGTTATTACAATGCTAA | |
| atlE_J_Rev | GTGGGATCCTTATTTTTTTAATGCTCGACTATCTACCCAGCC | |
| atlE_O_Fw | TCTGAGCTCAAGGAGGAGACTGACCATGAAAAAAATCATTTCAGGTATGTT | |
| atlE_O_Rev | GCGGGATCCTTAATTCACTTTTTGTACATAACGTT | |
| atlE_O_pETF | CCCTCTAGAAATACTTTTGTTTAACTTTAAGAAGGAGATATACGATGGAAGAGCTTGTAAAAACAGAAAC | |
| atlE_O_pETR | ATGGGATCCATTCACTTTTTGTACATAACGTTTATTTGAAGTGAT | |
| pTetH_Fw | GCTTGATCGTAGCGTTAACAGATCTACTC | |
| pTetH_Rev | CAAATTGTGGATGTGACCATGCGG | |
| pGhost_Fw | GTCACGACGTTGTAAAACGACGG | |
| pGhost_Rev | CTAGCGGACTCTAGAGGATCCCA | |

Erm$^{R}$, resistance to erythromycin

Amp$^{R}$, resistance to ampicillin
